# Cultivation of halophilic archaea in shallow subsurface martian conditions has implications for extant life on Mars

**DOI:** 10.64898/2026.07.11.737928

**Authors:** Adam Robinson, Shannon McQuaig-Ulrich, Tyler Dondero, Aaron Celestian, Scott M. Perl

**Affiliations:** Space Life Sciences Lab, University of Florida, 505 Odyssey Way, Merritt Island, FL 32953; Natural Sciences, St. Petersburg College, 2465 Drew St, Clearwater, FL 33765; Mineral Sciences, Los Angeles Natural History Museum, 900 Exposition Blvd, Los Angeles, CA 90007; Blue Marble Space Institute of Science, 600 1st Avenue, Seattle, Washington 98104

## Abstract

The present-day martian surface is generally considered inhospitable to life because of low atmospheric pressure, intense surface radiation, global desiccation, and oxidizing chemistry which has been increasing since the late Noachian. However, shallow martian subsurface regions where mineralogy has shown groundwater movement may include localized hypersaline environments capable of retaining liquid water and supporting microbial metabolism. *Haloferax volcanii*, a model halophilic archaeon, has previously been shown to survive under low-pressure martian conditions (24 mbar) and to grow anaerobically supported by the Mars-relevant oxyanions nitrate and perchlorate under high-salinity conditions. Here, we investigated whether *H. volcanii* could actively grow under a combination of environmental and chemical conditions relevant to potentially habitable shallow subsurface martian lacustrine settings. Cultures were incubated for 160 days under anoxic, CO₂-enriched, low-pressure conditions (24 mbar) in hypersaline liquid media supplemented with nitrate or perchlorate. Growth was observed in all low-pressure treatments and was confirmed by increases in optical density and biological reduction of nitrate and perchlorate. Scanning electron microscopy revealed extensive biofilm formation in low-pressure cultures, and Raman spectroscopy demonstrated the persistence of carotenoid biosignatures after prolonged incubation under martian conditions. Water loss remained below 4% across all treatments, indicating long-term stability of hypersaline brines throughout the experiment. These results demonstrate for the first time that a halophilic archaeon is capable of active growth and metabolism under a Mars-relevant combination of low pressure, high salinity, anoxia, and oxidizing chemistry, providing experimental support for the potential habitability of localized shallow subsurface martian environments.

**Importance:** The search for cellular life is a major objective of future Mars exploration. While many studies have examined whether microorganisms can survive under martian conditions, far fewer have demonstrated active growth and metabolism. Here, we document *Haloferax volcanii* as the first halophilic archaeon capable of active growth under a defined combination of Mars-relevant low atmospheric pressure, high salinity, anoxia, and oxidizing chemical conditions. These findings expand the current understanding of the environmental limits of microbial growth and provide experimental evidence that localized brine environments in the shallow martian subsurface could support active microbial metabolism, if suitable organics and liquid water are present. In addition, this study establishes a practical framework for cultivating halophilic microorganisms under low-pressure martian conditions and may help guide future efforts to detect, cultivate, and characterize potential extant life on Mars.

## Introduction

Humankind’s first extraterrestrial life-detection experiment was conducted on the surface of Mars in 1976 during NASA’s Viking mission. Specifically, the Labeled Release (LR) experiment aboard the Viking 1 lander was designed to inoculate martian regolith with a radiolabeled nutrient solution and detect the evolution of gaseous metabolic byproducts as evidence of active microbial metabolism (1). The instrumentation recorded an initial “positive” response, which the experiment’s investigators interpreted as being consistent with biological activity on the martian surface (2,3). While the Viking biology experiments were a technical and operational success, the interpretation of the LR results has remained controversial (4), and the prevailing consensus is that the experiment likely did not detect extant life on the martian surface (5). In the five decades since Viking, substantial advances in our understanding of the martian environment, including low atmospheric pressure, intense UV-C radiation, extreme desiccation, the widespread presence of (per)chlorates, and a lack of stable liquid water, suggest that the present-day martian surface is highly biocidal and unlikely to support extant life (6,7,8,9). However, a growing body of work suggests that the martian subsurface may remain conducive to extant life, owing to the potential for microenvironments with higher atmospheric pressure, shielding from surface radiation, and the potential for liquid water in the form of brines (10,11,12,13,14,15,16,17).

Efforts to constrain the habitability of present-day Mars have been driven by planetary protection and astrobiology research questions (18,19,20). Best efforts to decontaminate spacecraft and landers sent to Mars from Earth are often incomplete as many microbial isolates have been recovered from spacecraft assembly clean rooms (21,22,23,24,25). Planetary protection efforts that seek to understand the risk for forward contamination on Mars have shown that numerous bacterial species, including isolates from spacecraft clean rooms and the immediate surrounding areas, can survive or grow under low-pressure conditions relevant to the martian surface (8,26,27,28,29,30). Specifically, organisms that can grow at atmospheric pressures representative of the martian surface and shallow subsurface (7-10 mbar) are categorized as hypopiezotolerant (8,31). At shallow subsurface depths, microenvironments within fluid inclusions inside salt or ice may allow for higher pressures (25 mbar) at depths of at least ∼20 cm below the surface (7,8).

Furthermore, very shallow subsurface rocks (cm-to-mm) within meter-depth crater sites have shown a wide array of secondary pore spaces which would allow for pressures to be homogenous, and representative of deeper pressures should the surface crusts remain undisturbed (98). Pioneering low-pressure microbiology work expanded our understanding on the risks of contaminating potentially habitable regions on Mars with bacterial species present on spacecraft and landers sent from Earth (8). Moreover, recent work demonstrated that fungal isolates recovered from NASA spacecraft assembly cleanrooms can survive conditions relevant to the martian surface, including low pressure, CO₂-enriched atmospheres, and UV radiation (32). However, much less is known about the ability of fungi or archaea to grow under low pressure martian conditions (8).

The detection of methane on Mars (33,34) has driven speculation on the potential for extant methanogenic microorganisms in the subsurface of Mars (35,36). Methanogenic archaea have been used extensively as an astrobiology model for understanding the potential for extant life on Mars (36,37,38,39,40,41,42). However, many low-pressure and polyextreme combination Mars simulation experiments primarily look at survival under certain defined polyextreme martian conditions, and not growth or active metabolism. To date, the methanogen *Methanosarcina barkeri* is the only archaeal species demonstrated to be capable of active metabolism under defined low-pressure conditions representative of the shallow martian subsurface, although robust growth under these conditions has not been observed (36). Studies on the ability of extremophiles and polyextremophiles to survive martian conditions are replete in the literature, but studies on growth and active metabolism under low-pressure martian conditions are scarce, yet essential to constrain the habitability of present-day Mars (8). This may be due, in part, to the technical challenges associated with cultivating microorganisms under low-pressure martian conditions. Because the atmospheric pressure on Mars is at or below the vapor pressure of liquid water, growth media can undergo rapid evaporation, resulting in severe desiccation that can make sustained microbial cultivation extremely difficult (7,8). Hypersaline liquid media, by contrast, exhibit relative stability under Mars-relevant pressures over extended durations and demonstrate markedly reduced susceptibility to evaporative water loss compared with conventional microbial growth media. This stability facilitates long-duration cultivation experiments designed to evaluate microbial growth and metabolic activity under low-pressure martian conditions (43).

Halophilic microorganisms have been another important extremophilic model for astrobiology studies due to the existence or predicted existence of extraterrestrial brines throughout our solar system (44,45,46,47,48,49,50,51), and particularly with relevance to the shallow martian subsurface (43,52). Notably, the model halophilic archaea *Haloferax volcanii* has been the subject of several previous astrobiology studies (43,45,49,53,54). For example, the first attempt to cultivate halophilic archaea under defined shallow subsurface martian conditions documented the survival of *H. volcanii* for approximately four months but did not observe growth. However, higher survival rates and distinct morphological changes were observed under low-pressure martian conditions relative to Earth-atmosphere controls, suggesting a stress response to low-pressure environments (43). More recently, *H. volcanii* was shown to grow anaerobically supported by perchlorate, but only at salinities higher than those preferred during aerobic growth. Furthermore, *H. volcanii* tolerated and grew anaerobically at perchlorate concentrations exceeding those reported for any other microorganism to date, demonstrating an exceptional capacity to withstand the combined stresses of high salinity and perchlorate exposure (54). These studies have uncovered previously unknown metabolic requirements for anaerobic growth on Mars-relevant electron acceptors, highlighting the importance of salinity and electron acceptor availability in shaping microbial habitability under martian conditions, while guiding future efforts to understand active microbial metabolism under low-pressure martian conditions.

Here, we document *Haloferax volcanii* as the first halophilic archaeon capable of active growth under low-pressure conditions that simulate potentially habitable subsurface niches on Mars. This was achieved by incubating *H. volcanii* under low-pressure (24 mbar), anoxic (CO₂-enriched) atmospheric conditions in hypersaline liquid media containing the Mars-relevant oxyanions perchlorate or nitrate (97) as alternative electron acceptors. Additionally, this study used scanning electron microscopy (SEM) to observe potential morphological changes and Raman spectroscopy to investigate potential chemical changes in halophilic biosignatures (i.e. carotenoid pigmentation) due to the simulated martian conditions. The conditions chosen for this study balance the limits of the physiological capabilities of *H. volcanii* (43,54) while maintaining environmental conditions that are plausible in the shallow martian subsurface today (7,8). Most importantly, we establish a practical cultivation framework for growing halophilic microorganisms in liquid media under Mars-relevant environmental conditions, providing a foundation for future studies of microbial growth, metabolism, and biosignature preservation in simulated subsurface martian habitats.

## Results

### Growth of *H. volcanii* under low-pressure martian conditions (24-mbar)

Cultures supplemented with nitrate and perchlorate that were incubated for 160 days during the Mars simulation experiment showed statistically significant increases in optical density at 630 nm (Fig. 1A). Because dissolved-oxygen controls received the same starting inoculum and showed no significant increase in OD, increases in OD observed in nitrate- and perchlorate-supplemented cultures are unlikely to reflect carryover biomass alone, indicating active growth. However, growth under martian atmospheric pressure was significantly lower than that of control cultures incubated under standard Earth atmospheric pressure (Fig. 1B). At both Mars and Earth atmospheric pressures, anaerobic growth on nitrate was significantly greater than that on perchlorate (Figs. 1A, B). Semi-continuous monitoring of growth for Earth control cultures at ∼1013 mbar and 21°C showed a ∼6-8-week lag phase for anaerobic growth on 100 mM nitrate, a ∼7-8-week lag phase for 20 mM nitrate, a ∼11-week lag phase for 100 mM perchlorate, and a ∼5-week lag phase for 20 mM perchlorate (Fig. 2). The large standard deviations observed at certain time points in Fig. 2, particularly during the transition from lag to exponential growth, reflected asynchronous growth initiation among biological replicates. Variability subsequently decreased as all replicates entered active growth and exhibited similar growth trajectories.

**Figure 1:**
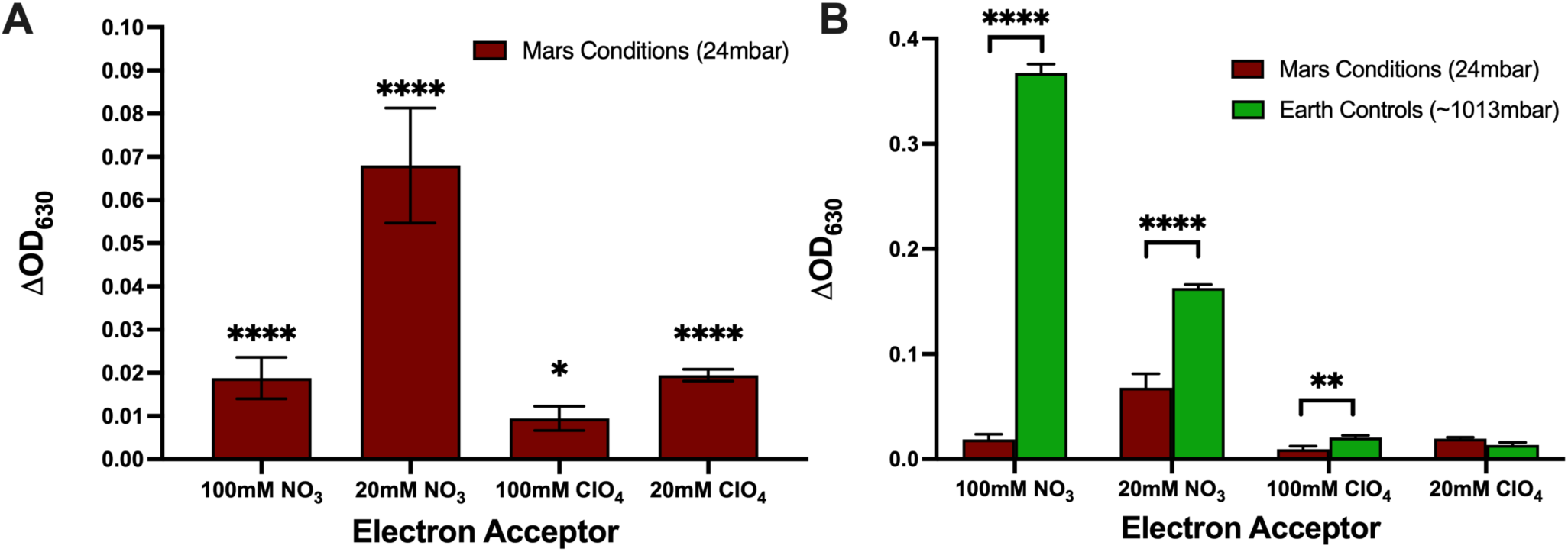
End-point optical density (OD_630_) measurements of Haloferax volcanii cultures following 160 days of incubation during the Mars simulation experiment. (**A**) OD_630_ increases above baseline for cultures incubated under simulated Martian atmospheric pressure (24 mbar). (**B**) Comparison of end-point OD_630_ values between cultures incubated under simulated Martian atmospheric pressure (24 mbar) and Earth control cultures incubated under identical chemical conditions at standard Earth atmospheric pressure (∼1013 mbar). Bars represent mean ± SD (n = 3), and asterisks indicate statistical significance, * p < 0.05, ** p < 0.01, **** p < 0.0001.

**Figure 2:**
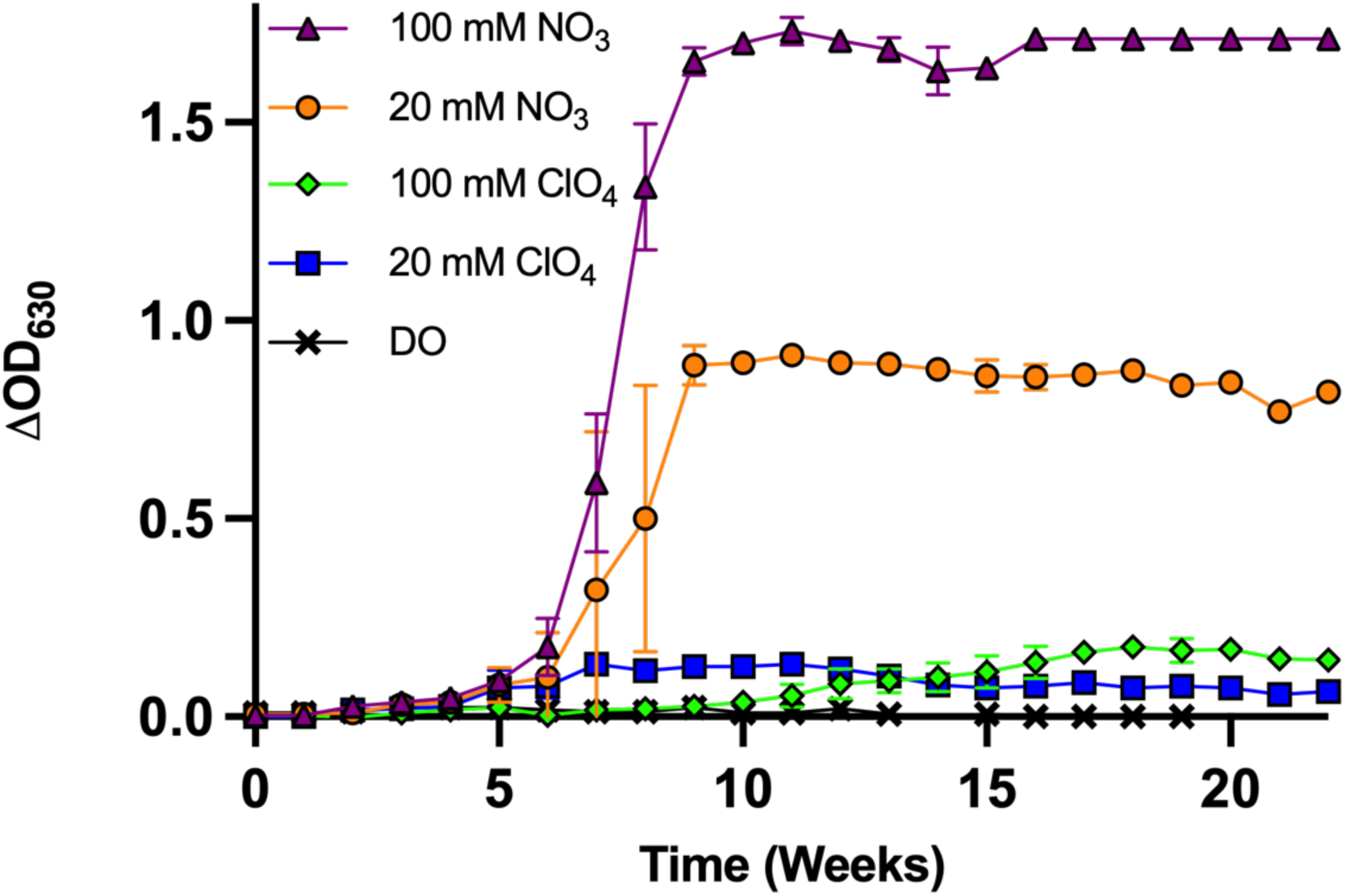
Semi-continuous monitoring of anaerobic growth by H. volcanii during incubation under Earth control conditions (∼1013 mbar, 21°C) over 160 days. Growth was monitored by optical density measurements (OD_630_) for cultures supplemented with 100 mM nitrate, 20 mM nitrate, 100 mM perchlorate, 20 mM perchlorate, or no added electron acceptor (control dissolved oxygen, DO). Data points represent mean ± SD (n = 3).

### Viability Following Long-Term Incubation Under Mars Conditions

Fresh cultures inoculated from aliquots of cultures incubated for 160 days under Mars simulation conditions showed increases in optical density (OD_630_), indicating active growth and confirming cell viability across all Mars simulation and Earth control treatments (Figs. 3A, B). Furthermore, when viability assay cultures were incubated under oxygenated conditions at 37°C, significantly greater growth was observed in standard ATCC 974 medium containing 125 g L⁻¹ NaCl compared to modified medium containing 225 g L⁻¹ NaCl (Figs. 3A, B).

**Figure 3:**
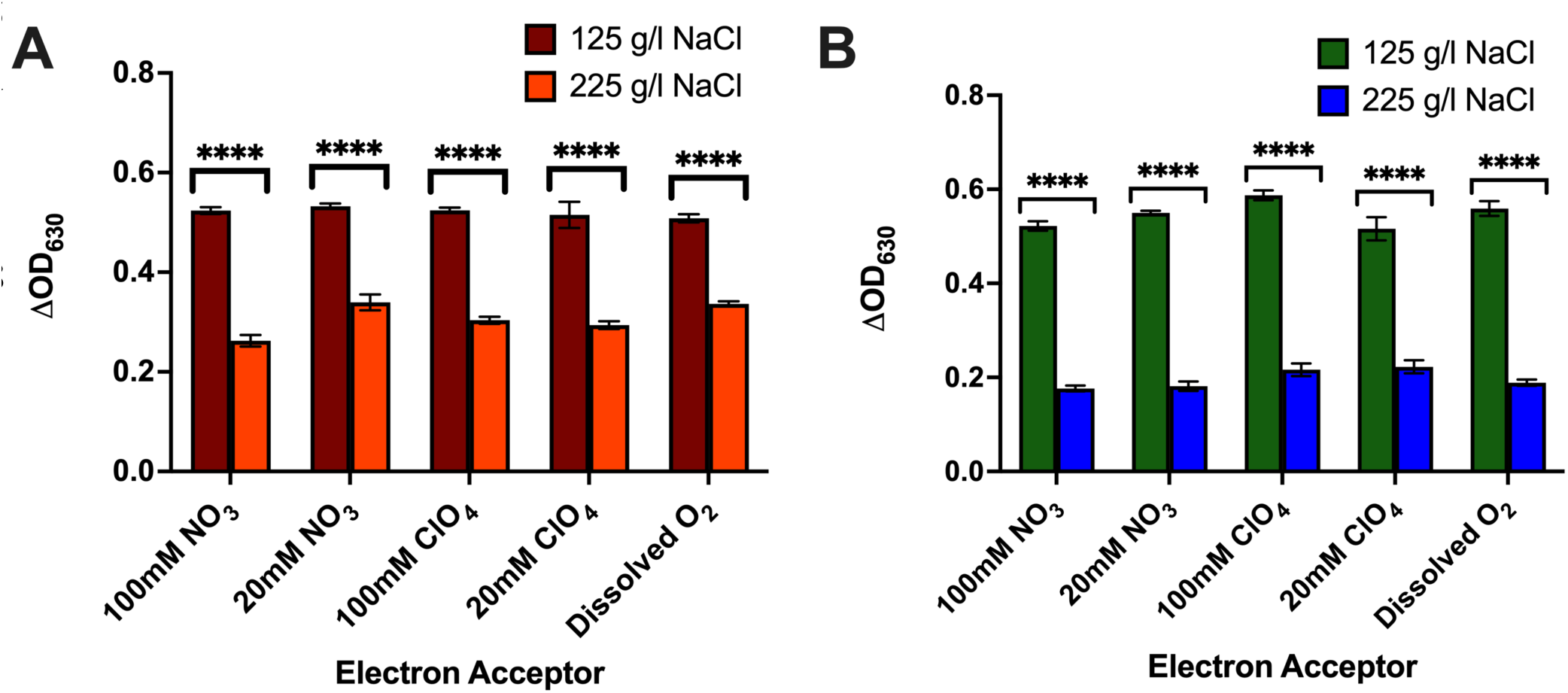
Optical density (OD_630_) measurements of viability assay cultures incubated under oxygenated conditions for 7 days at 37°C, following the completion of the Mars simulation experiment. 100 µL aliquots from each Mars simulation culture were used to inoculate triplicate tubes with 5 ml of ATCC 974 media containing either 125 g/L or 225 g/L NaCl. **A**) Viability assay of cultures incubated for 160 days under 24 mbar “Mars” conditions. **B**) Viability assay of control cultures incubated for 160 days at ∼1013 mbar Earth conditions. Bars represent mean ± SD (n = 3), and asterisks indicate statistical significance, **** p < 0.0001.

### Stability of Hypersaline Liquid Growth Media at 24 mbar

After 160 days of incubation under shallow subsurface martian atmospheric conditions, total water loss from 10 mL cultures remained minimal across all treatments (Fig. 4). Cultures containing 100 mM and 20 mM nitrate exhibited 2.13% and 2.05% water loss, respectively, while 100 mM and 20 mM perchlorate treatments showed 2.02% and 1.55% water loss. Cultures containing dissolved oxygen experienced the greatest water loss (3.76%), followed by media-only blanks (2.88%). Notably, water loss remained below 4% in all treatments, demonstrating long-term brine stability under low-pressure martian conditions.

**Figure 4:**
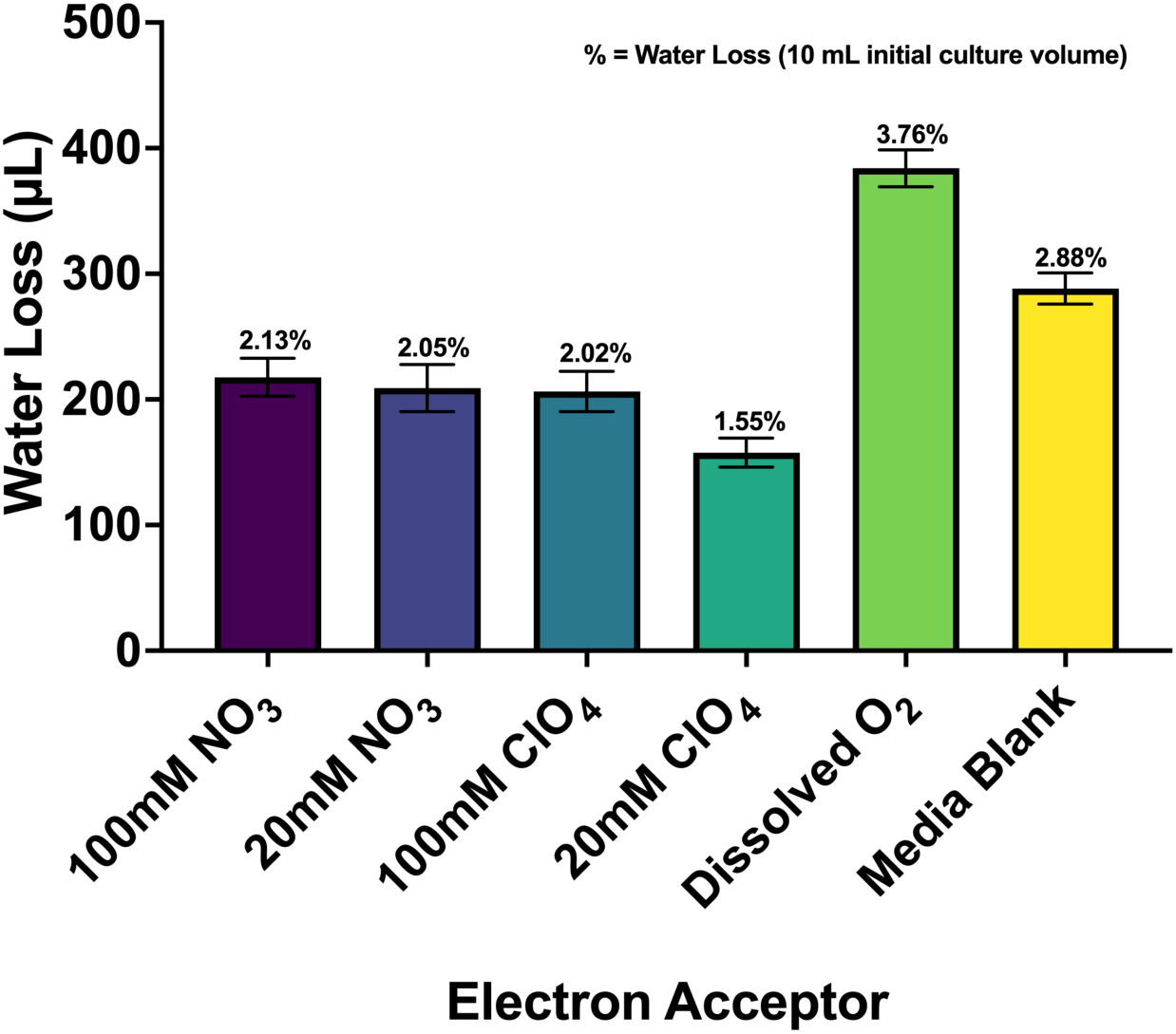
Water loss (µL) from 10 mL cultures incubated after 160 days of exposure to shallow subsurface martian atmospheric pressures (24 mbar). Percentages indicate total water loss relative to the initial culture volume. Across all electron acceptor treatments and media-only controls, water loss remained below 4%, demonstrating long-term brine stability under low-pressure martian conditions. Bars represent mean ± SD (n = 3).

### Morphological Changes Associated with Growth Under Mars Atmospheric Conditions

After 160 days of incubation, SEM analysis revealed pressure-dependent changes in cellular organization and surface morphology (Fig. 5). Specifically, under Earth control conditions, cells generally retained morphology similar to that of healthy preculture populations, particularly at 20 mM nitrate and perchlorate (Figs. 5C, G). However, higher concentrations (100 mM) were associated with visible cell distortion (Figs. 5A, E). In contrast, cultures incubated under low-pressure martian conditions exhibited substantial structural reorganization, characterized by dense cellular aggregation and the development of biofilm-like structures with cells embedded in an apparent extracellular matrix. This dense biofilm material was not observed in Earth control cultures or low-pressure baseline controls that did not demonstrate detectable growth, suggesting that the formation was associated with active growth under martian pressure. Higher-magnification (x12,000) visualization revealed cells embedded in the dense biofilm matrix and cellular appendages extending outward (Fig. 6).

**Figure 5:**
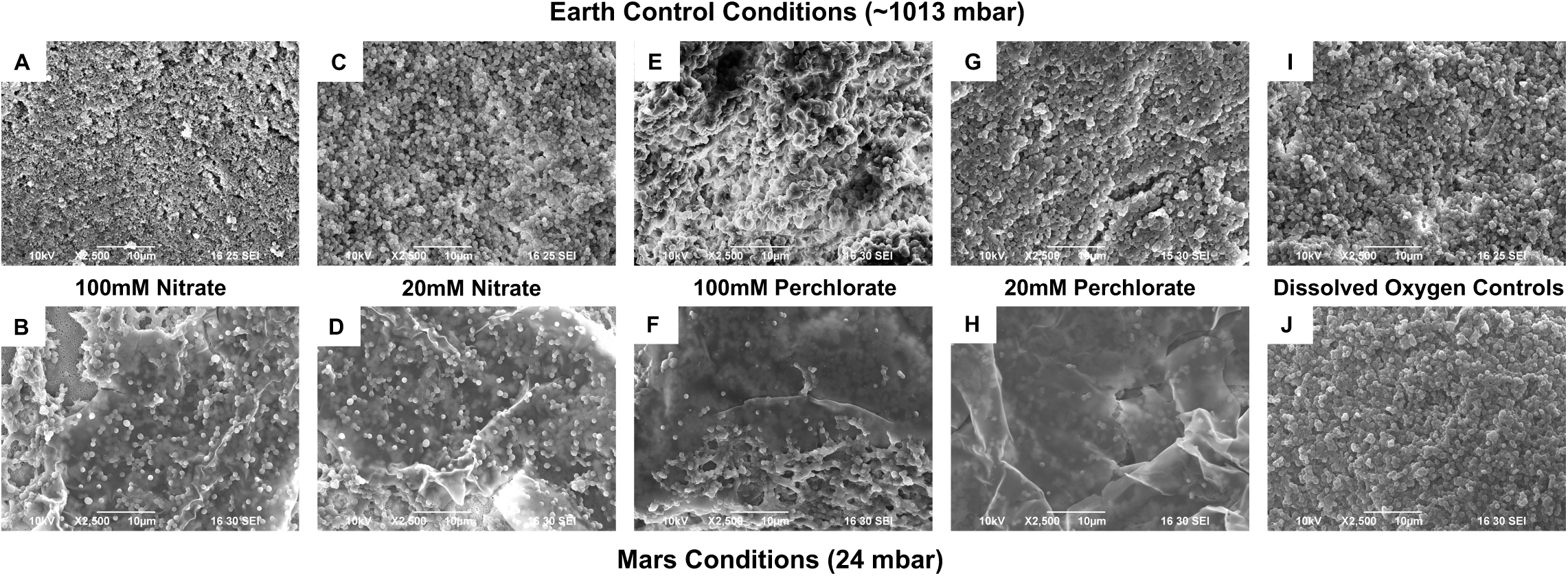
Scanning electron microscopy (SEM) micrographs (2500x magnification) of Haloferax volcanii after 160 days of incubation under Earth control conditions (∼1013 mbar; top row, A,C,E,G,I) and martian pressure conditions (24 mbar; bottom row, B,D,F,H,J). Baseline controls without added alternative electron acceptors displayed a morphology mostly consistent with that of healthy preculture cells under Earth (Fig. 5I) and Mars (Fig. 5J) conditions. Earth control cultures grown anaerobically with 20 mM nitrate or perchlorate (Figs. 5C, G) exhibited morphology similar to preculture cells, whereas cells grown with 100 mM nitrate or perchlorate (Figs. 5A, E) showed evidence of cell distortion. In contrast, cultures incubated under low-pressure martian conditions (Figs. 5B,D,F,H) displayed pronounced biofilm formation, with cells embedded within an extracellular biofilm matrix. Notably, this matrix was absent in low-pressure baseline controls that did not exhibit detectable growth (Fig. 5J). Scale bars = 10 µm.

**Figure 6:**
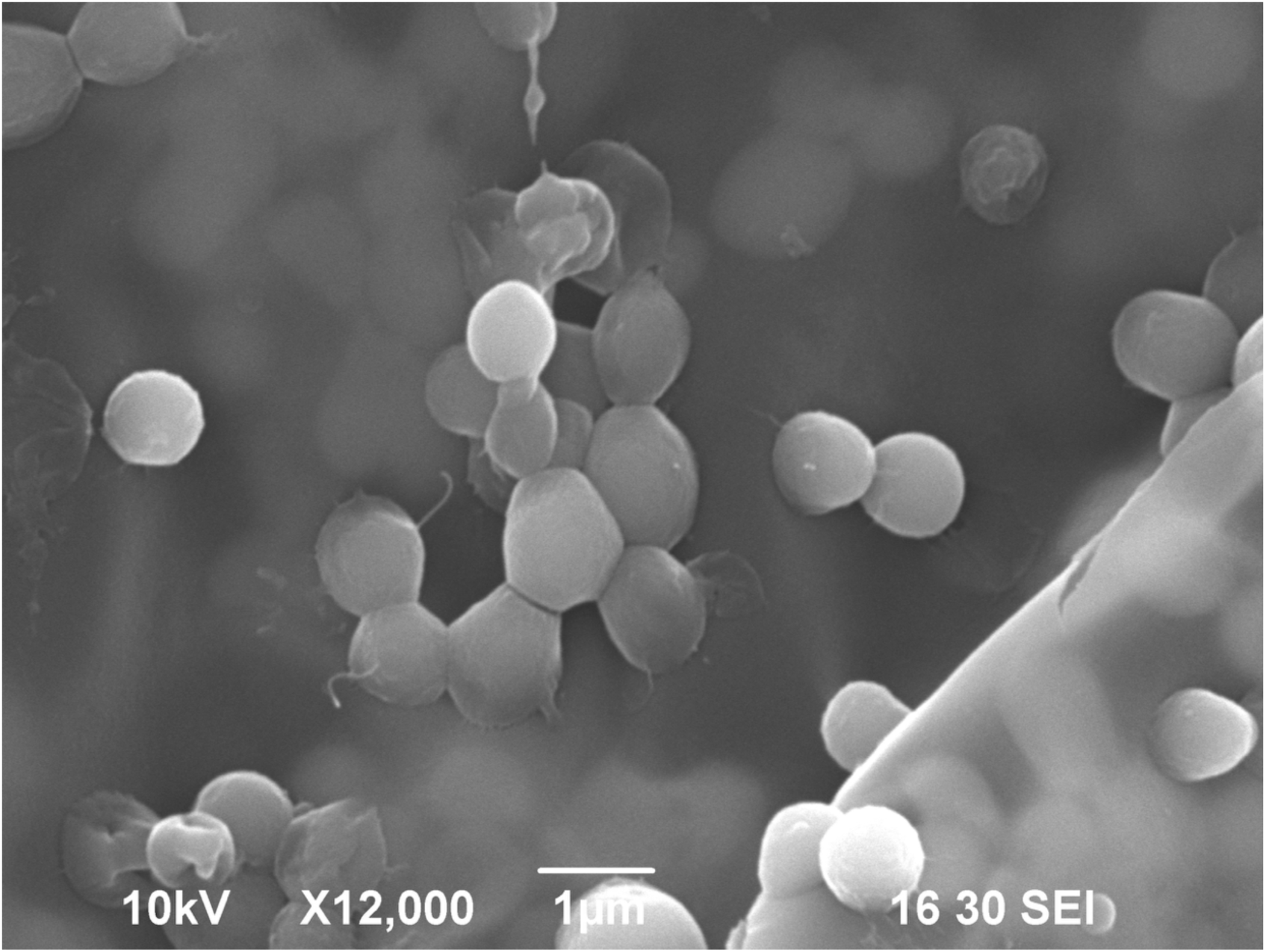
Scanning electron microscopy (SEM) micrograph (12000x magnification) of H. volcanii grown with 100 mM nitrate at 24 mbar. Higher-magnification imaging provides a detailed view of cells embedded within the biofilm matrix. Scale bars = 1 µm.

### Persistence of Raman Carotenoid Biosignatures

Spectral peaks corresponding to carotenoid biosignatures (∼1505, ∼1150, and ∼1000 cm⁻¹) were clearly visible in cultures incubated under dissolved oxygen control conditions (Fig. 7). These peaks were also detected in cultures grown anaerobically supported by nitrate or perchlorate, indicating the continued presence of carotenoid pigments following anaerobic growth under both martian and Earth atmospheric pressures. However, carotenoid peak intensities were reduced in anaerobic cultures relative to dissolved oxygen controls. Under simulated martian atmospheric pressure (24 mbar), nitrate-supported cultures generally exhibited stronger carotenoid peaks than perchlorate-supported cultures. This may be in part due to the accumulation of chlorite resulting from partial perchlorate reduction, which has been previously suggested to degrade carotenoid biosignatures (54). In contrast, carotenoid peaks in Earth-pressure cultures remained detectable but were substantially reduced relative to dissolved oxygen controls, with less pronounced differences between nitrate- and perchlorate-supported growth (Fig. 7B).

**Figure 7:**
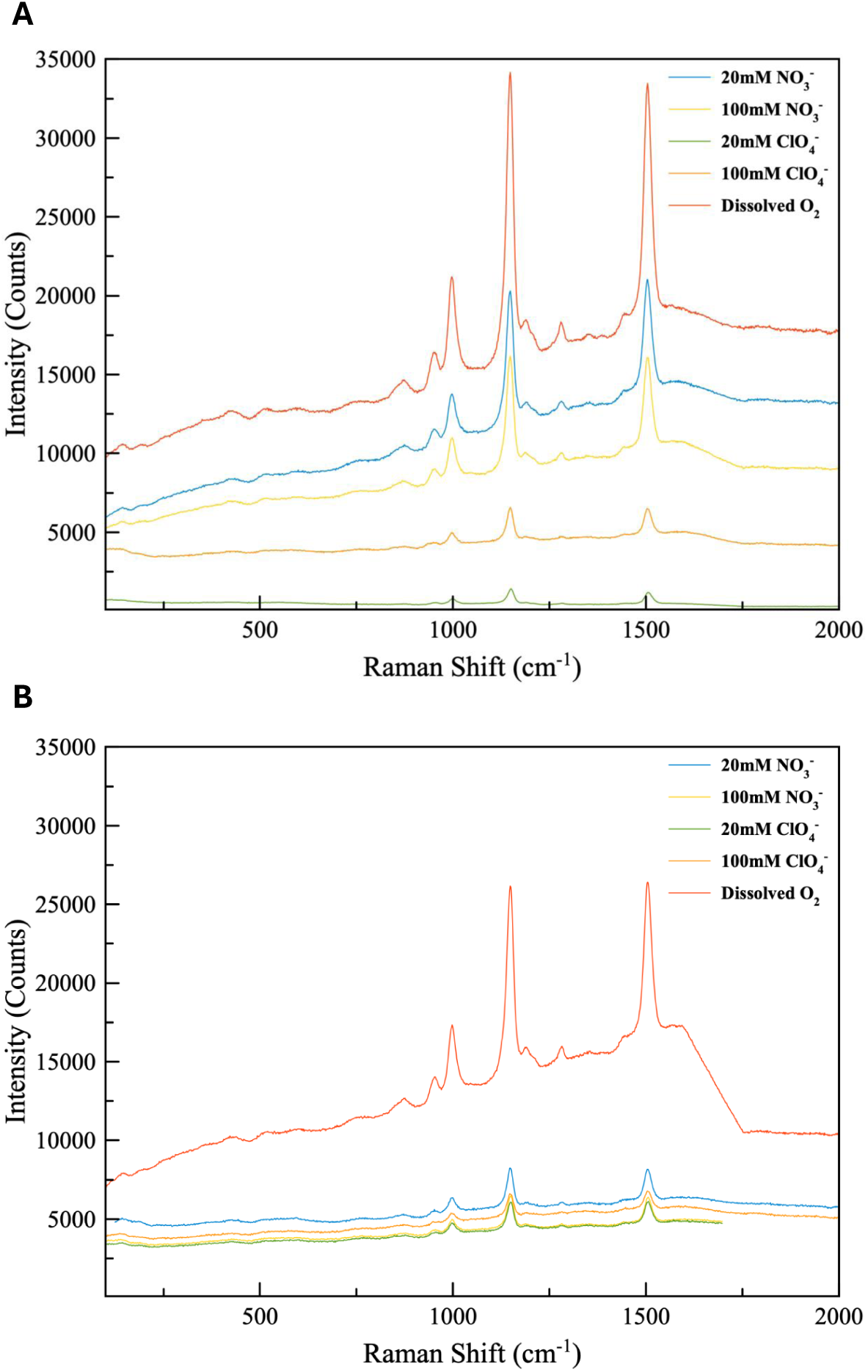
**A**) Raman spectra of cultures incubated in low-pressure martian conditions. **B**) Raman spectra of control cultures incubated at standard Earth atmospheric conditions. Raman peaks associated with halophilic carotenoids are slightly diminished due to the presence of nitrate and perchlorate.

### Biological Reduction of Mars-Relevant Oxyanions During Anaerobic Growth

Significant decreases in nitrate and perchlorate concentrations were observed following 160 days of incubation under both simulated martian (24 mbar) and Earth atmospheric pressures (∼1013 mbar) (Fig. 8). In nitrate-supplemented cultures, nitrate concentrations decreased by ∼29% and ∼33% in the 20 mM Mars and Earth treatments, respectively, and ∼15% and ∼44% in the 100 mM Mars and Earth treatments (Fig. 8A). Similarly, perchlorate concentrations decreased by ∼29% and ∼28% in the 20 mM Mars and Earth treatments, respectively, and ∼16% and ∼21% in the 100 mM Mars and Earth treatments (Fig. 8B). These results indicate utilization of nitrate and perchlorate during incubation and are consistent with the observed increases in optical density (Fig. 1) suggesting growth of cultures incubated under simulated martian conditions.

**Figure 8.**
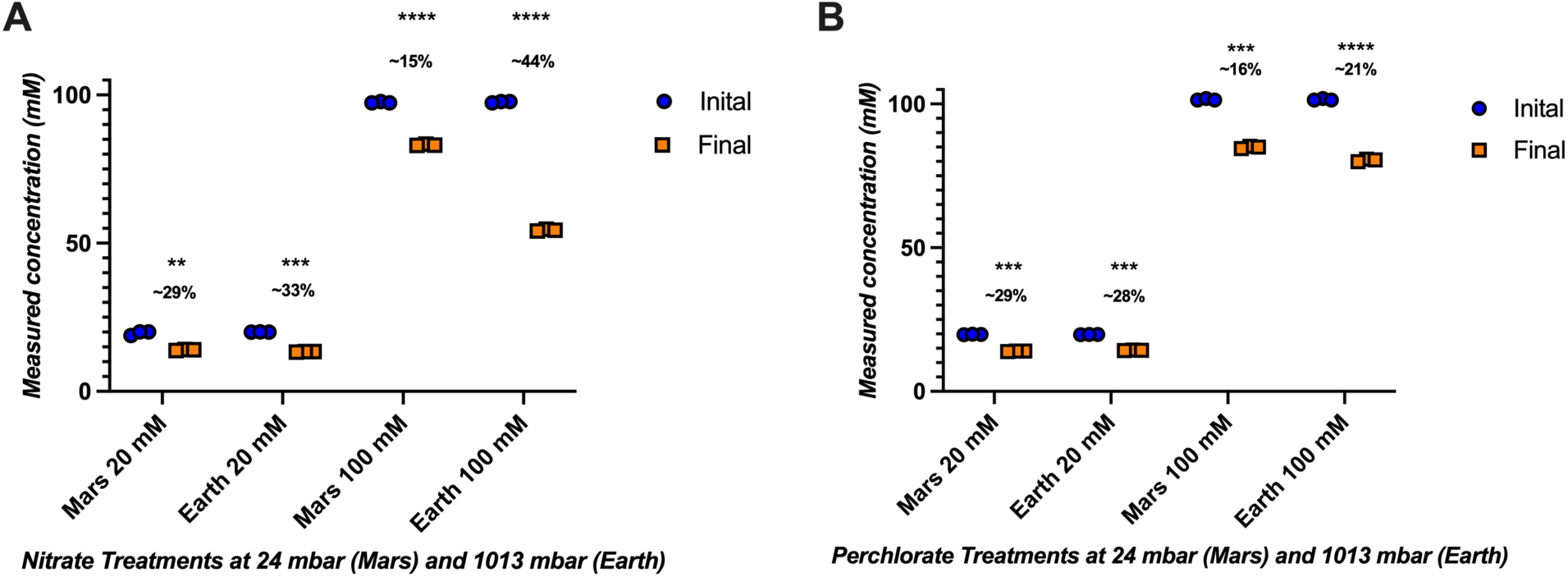
Reduction of nitrate and perchlorate during 160-day incubation under simulated martian and control Earth atmospheric pressures. (A) Initial and final nitrate concentrations in cultures supplemented with 20 or 100 mM nitrate and incubated at 24 mbar (Mars) or ∼1013 mbar (Earth). (B) Initial and final perchlorate concentrations in cultures supplemented with 20 or 100 mM perchlorate and incubated under the same conditions. Points represent individual biological replicates (n = 3). Percent values indicate mean reduction relative to initial concentrations. Statistical significance was determined using two-tailed paired t-tests comparing initial and final concentrations within each treatment (*p < 0.05, **p < 0.01, ***p < 0.001, ****p < 0.0001).

## Discussion

### Growth And Brine Stability Under Martian Atmospheric Conditions

We have shown that *H. volcanii* can grow under anoxic, CO₂-enriched, low-pressure (24 mbar) conditions representative of a hypothetical habitable hypersaline niche within a sealed fluid inclusion in a martian salt mineral at depths of at least 20 cm below the surface. This is supported by statistically significant increases in optical density at 630 nm indicating growth (Fig. 1A), coupled with depletion of the supplied oxyanion electron acceptors (Fig. 8). However, growth was significantly lower under low-pressure martian conditions (24 mbar) than in control cultures incubated under anoxic conditions at standard Earth atmospheric pressure (∼1013 mbar), highlighting the inhibitory effects of hypobaria on anaerobic growth. (Fig. 1B). While growth was observed in all tested conditions, the effects of lower temperatures on anaerobic growth were evident, with lag phases of ∼5-11 weeks in the Earth atmospheric pressure controls (Fig. 2), compared to the 3–4 day lag phase previously reported during anaerobic growth at a preferred temperature of 37°C (54). Growth at 24 mbar contrasts with Robinson and McQuaig-Ulrich (43), who demonstrated that *H. volcanii* could survive under similar low-pressure martian conditions for up to 113 days but did not observe growth. Notably, that study used a growth medium containing the optimal NaCl concentration for aerobic growth (125 g L⁻¹); however, anaerobic growth using alternative electron acceptors was not observed, even under Earth control conditions (43). Subsequent work demonstrated that the optimal conditions for aerobic and anaerobic growth in *H. volcanii* may differ substantially. For example, anaerobic growth supported by perchlorate in *H. volcanii* is only possible when NaCl concentrations are increased to 175 g L⁻¹, while anaerobic growth in the presence of 200 mM perchlorate required NaCl concentrations of 225 g L^−1^ (54). By addressing critical gaps in our understanding of the physiological requirements for anaerobic growth in this model halophile, here we were able to successfully cultivate a halophilic microorganism under low-pressure martian conditions for the first time.

In addition to supporting anaerobic growth, the elevated salt concentration helped stabilize the liquid growth medium under conditions that would otherwise promote rapid evaporation. This likely contributed to the long-term persistence of liquid water throughout the 160-day incubation period at 24 mbar and enabled sustained microbial growth under Mars-relevant pressure conditions. In Robinson and McQuaig-Ulrich (43), a hypersaline growth medium containing 125 g L^−1^ NaCl was incubated for 113 days at 24 mbar and 21 °C, with maximum water loss of ∼15%. In contrast, the growth medium used in the present study contained 225 g L^−1^ NaCl and exhibited a maximum water loss of only 3.76% after 160 days. Interestingly, lower levels of water loss were observed in all samples where active microbial growth occurred (1.55–2.13%) compared to uninoculated controls or treatments in which no growth was detected (Fig. 4).

Macroscopically, cultures incubated under low-pressure conditions exhibited visible biomass sediment accumulation, turbidity throughout the water column, and a surface film at the air–liquid interface. Microscopic examination revealed dense biofilm formation exclusively in low-pressure treatments that supported microbial growth (Figs. 5 and 6). Together, these observations suggest that, in addition to the intrinsic water-retaining properties of the hypersaline medium, biofilm formation at the air–liquid interface may have further reduced evaporative water loss and contributed to the long-term stability of the cultures. These findings suggest that hypersaline conditions not only support anaerobic growth under low-pressure martian atmospheric conditions but also promote the long-term stability of liquid water necessary to sustain active microbial metabolism.

### Morphology and Biosignature Alteration

Morphological changes associated with stress conditions have been previously documented in halophilic archaea (43,62,63,64,65,66,67). For example, low water activity has been reported to trigger cell size reduction in *Halobacterium salinarum* (63), while exposure to perchlorate can cause cellular distortion at elevated concentrations (45). In *H. volcanii*, mechanical compression has been shown to induce morphological transitions resembling multicellular tissue-like structures (66). Under low-pressure conditions, *H. volcanii* has also been reported to exhibit alterations in cell morphology following survival for 113 days at 24 mbar, including changes in cell size, the formation of cellular appendages, and biofilm production (43). Similarly, growth of the bacterium *Serratia liquefaciens* under simulated martian conditions (7 mbar) resulted in pronounced ultrastructural changes, including cell distortion, thinning of the cell wall, loss of fimbriae, and evidence of disrupted cell division (68). In the present study, slight cellular distortion was observed in cultures grown anaerobically at Earth atmospheric pressure, although cells generally retained their typical morphology (Figs. 5A, C, E, G, I). In contrast, the most pronounced morphological changes were observed in cultures grown under low-pressure martian conditions. Dense biofilm formation was observed exclusively in low-pressure treatments (24 mbar) that supported active growth (Figs. 5B, D, F, H), with individual cells and cellular aggregates embedded throughout the biofilm matrix (Figs. 6,7). Control cultures incubated for the same duration at 24 mbar that did not exhibit active growth, displayed only minor cellular distortion (Fig. 5J) and lacked the extensive biofilm structures observed in the actively growing cultures. Taken together, these observations suggest that biofilm formation was associated with active growth under low-pressure martian conditions and may represent a physiological response that enhances survival and persistence under hypobaric stress.

Biological pigments have been proposed as potential biosignatures in the search for life beyond Earth (69,70). In particular, carotenoid pigments produced by halophilic microorganisms generate distinct Raman spectral peaks that can serve as indicators of halophilic life (61). The carotenoids β-carotene and bacterioruberin, share Raman spectral peaks based on their molecular structure and vibrational sensitivity, and allow for a proxy for cell growth in extant settings as well as a biomarker after cell death (51). Consequently, halophile-derived carotenoid signatures have been suggested as potential targets in the search for evidence of life on Mars (51,71,72,73). However, the preservation of these biosignatures may be influenced by environmental stressors relevant to the martian environment. For example, β-carotene exposed to simulated martian surface conditions for 722 days outside the International Space Station exhibited significant degradation when analyzed using Raman spectroscopy (74). Similarly, previous work demonstrated that when *H. volcanii* is grown anaerobically supported by perchlorate at 37 °C, Raman peaks associated with the carotenoid bacterioruberin are severely diminished and difficult to detect (54). In the present study, Raman peaks associated with bacterioruberin remained detectable in cultures that exhibited active growth under low-pressure martian conditions, although peak intensities were somewhat reduced relative to controls in which anaerobic growth did not occur (Fig. 7). The persistence of detectable bacterioruberin signatures contrasts with the observations of Robinson et al. (54), where carotenoid peaks were substantially diminished during anaerobic growth on perchlorate. One notable difference between the studies was incubation temperature (21 °C versus 37 °C), suggesting that lower temperatures may enhance the preservation of carotenoid biosignatures. If hypothetical martian microorganisms produce pigments analogous to carotenoids, such compounds may be preferentially preserved within the subsurface, where protection from surface UV-C radiation could promote long-term biosignature stability.

### Astrobiology in the Age of Humans to Mars

The National Academies Science Strategy for the Human Exploration of Mars has identified the search for life as a primary scientific objective of human exploration of Mars (78). Considerable effort has been devoted to the development of technologies capable of detecting biosignatures and signs of extinct or extant life on Mars (79,80,81,82,83,84,85,86). However, substantial gaps remain in the development of protocols designed to cultivate and physiologically characterize potential martian microorganisms following their detection. In many respects, a modern “Viking 2.0” cultivation strategy has yet to be fully developed (87). The original Viking Labeled Release experiment employed a nutrient solution (1) that would likely be unsuitable for many true halophilic microorganisms on Earth (88), illustrating the challenges associated with designing cultivation systems for unfamiliar forms of life. We should not be in a position where extant life is detected on Mars without a clear framework for how to proceed. Preparation for the potential discovery of martian life should therefore include strategies not only for detection, but also for containment, cultivation, and physiological characterization. Although martian microorganisms may ultimately resemble extremophiles on Earth that are difficult or impossible to culture under laboratory conditions (89,90), cultivation attempts should nevertheless be pursued in order to investigate their physiology, metabolism, environmental tolerances, and potential ecological roles and other biological characteristics that cannot be fully determined through culture-independent approaches alone (91). In the future, organisms such as *H. volcanii* and other halophilic, psychrophilic, or methanogenic microorganisms may serve as useful positive controls in cultivation assays designed to recover extant martian life, and the protocols outlined herein may provide a framework for their development.

Furthermore, current planetary protection policies for backward contamination suggest that any material reasonably suspected of containing extraterrestrial biology be handled under the highest levels of biological containment until it can be adequately analyzed (94,95,96). Consequently, the immediate question is not whether putative martian organisms are likely to be pathogenic to humans, plants, or animals, but whether extant life exists in regions that may be accessed by future robotic or human explorers. The potential presence of martian biology, and our ability to safely study it under high-containment conditions, represents a critical challenge for future exploration (95). In terrestrial high-containment laboratories (biosafety level 3 and 4), aerosolization represents one of the primary biosafety hazards because airborne particles can facilitate containment failure and potentially lethal human infections by pathogens such as *Bacillus anthracis* and Ebola virus (75). Similar concerns may arise during human exploration of Mars, where the low ambient surface pressure (76,77) could promote rapid boiling, evaporation, and aerosol generation from liquid-bearing subsurface samples. Furthermore, cultivation attempts involving extant martian life would likely occur under medium- to high-vacuum conditions (8), where rapid boiling and evaporation of liquid samples could facilitate the aerosolization and dispersal of martian biology. Accordingly, astrobiodefense and planetary protection efforts should focus not only on preventing environmental contamination, but also on improving our understanding of how life can grow under martian conditions and how potential martian organisms could be safely cultivated and studied under high vacuum within containment systems analogous to biosafety level 3 and 4 laboratories on Earth.

### Conclusion

We have shown that *H. volcanii* can actively grow under environmental and chemical conditions relevant to the present-day shallow martian subsurface. However, this study was not designed to replicate the full thermal variability of the martian surface or shallow subsurface, including diurnal and seasonal freeze–thaw cycles, nor the complete suite of chemical conditions presently understood to exist on Mars. Rather, the objective was to determine whether active metabolism is physically possible under Mars-relevant combinations of low atmospheric pressure, high salinity, and oxidizing chemistry, provided that liquid water is available in the form of brines. Moreover, the media used in many previous low-pressure Mars simulation studies, including those employed herein, are supplemented with carbon sources and nutrients that are not representative of the current understanding of biologically available organic compounds on Mars (87). Our future studies will investigate the capacity of halophilic and psychrophilic microorganisms to grow under conditions that more closely approximate the Amazonian/present-day martian environment by incorporating freeze–thaw cycles, defined Mars media (99) containing Mars-relevant organic compounds (e.g., acetate) (100), and Mars regolith simulants. Importantly, while average surface and shallow subsurface temperatures on Mars remain below the freezing point of water, this does not preclude the possibility of active metabolism or extant life. Notably, the psychrophilic haloarchaeon *Halorubrum lacusprofundi* was originally isolated from Deep Lake, Antarctica, where environmental temperatures can reach as low as −20°C (92). Although *Hrr. lacusprofundi* survives prolonged freezing conditions, active metabolism occurs at warmer temperatures of −1°C or higher (93). Furthermore, under laboratory conditions, *Hrr. lacusprofundi* exhibits optimal growth at approximately 30°C (92), illustrating that microorganisms adapted to cold, oligotrophic environments may nevertheless possess substantially higher growth optima when cultivated under favorable conditions. This suggests that the low average temperatures on Mars should not be viewed as an absolute barrier to extant microbial activity, provided that transient liquid water and suitable chemical energy sources are available. The results presented here do not imply that microbial life exists on Mars today.

However, they add to a growing body of evidence suggesting that the shallow subsurface of Mars may retain environmental conditions capable of supporting active metabolism and extant life (87,101,102,103).

## Materials and Methods

### Growth Media

*Haloferax volcanii (*Mullakhanbhai and Larsen) Torreblanca et al. (55) (ATCC 29605) stock cultures were grown aerobically at 37 °C in a shaker using modified ATCC-974 liquid medium containing 125 g L^−1^ NaCl, 50 g L^−1^ MgCl₂·6H₂O, 5 g L^−1^ K₂SO₄, 0.2 g L^−1^ CaCl₂·2H₂O, 5 g L^−1^ casamino acids, and 5 g L^−1^ yeast extract. Trace element and chelated iron solutions were prepared as described by Hattori et al. (56). The trace element solution contained 0.1 g L^−1^ Na₂MoO₄·2H₂O, 0.2 g L^−1^ MnCl₂·6H₂O, 2 mg L^−1^ CoCl₂·6H₂O, 0.1 g L^−1^ ZnSO₄·7H₂O, and 0.1 g L^−1^ CuSO₄·5H₂O. The chelated iron solution contained 0.1 g L⁻¹ FeSO₄·7H₂O and 0.1 g L^−1^ EDTA. One milliliter of each solution was added per liter of medium. The pH was adjusted to 7.0 using 1.0 M NaOH, and the medium was sterilized by autoclaving for 30 minutes (121°C, 15 psi).

Modified ATCC-974 medium containing 225 g L^−1^ NaCl and all aforementioned components was used for cultures incubated under simulated shallow martian subsurface atmospheric conditions (24 mbar, CO₂-enriched). Media were supplemented with either 20 mM or 100 mM of the Mars-relevant alternative electron acceptors nitrate or perchlorate. Stock solutions of NaNO₃ and NaClO₄ were filter sterilized using 0.22 µm hydrophilic polyvinylidene fluoride (PVDF) luer-lock syringe filters and added aseptically to sterile media to achieve the desired final concentrations. For martian-condition cultures, 10 mL of 225 g L^−1^ NaCl modified ATCC-974 medium was dispensed into sterile flat-bottom test tubes, followed by inoculation with 200 µL of stock culture. “Earth control” cultures, i.e., those incubated anaerobically at 1013 mbar, were prepared in sterile Hungate tubes containing 15 mL of sterile 225 g L^−1^ NaCl ATCC 974 medium and inoculated with 300 µL of preculture. Under anoxic conditions, *H. volcanii* depletes dissolved oxygen present in the medium and can utilize alternative electron acceptors when available; therefore, chemical reduction of the medium to remove dissolved oxygen was not required (57). Tubes lacking supplemented alternative electron acceptors served as negative controls to quantify growth supported solely by dissolved oxygen. All treatments were prepared in triplicate.

### Low-Pressure Cultivation Conditions

An incubation temperature of 21 °C was selected to permit anaerobic growth of *H. volcanii* under the experimental chemical conditions (43,54), while maintaining a low-pressure atmosphere (24 mbar CO₂) that may be possible within sealed fluid inclusions in salt minerals or ice in the shallow martian subsurface at depths of approximately 20 cm (7,8). Furthermore, 21 °C is within the range of transient temperatures that may occur during seasonal freeze–thaw cycles and warm summer periods in equatorial regions of Mars (58,59). The experimental design was adapted from the protocols described in Robinson and McQuaig-Ulrich (43), which was originally inspired by Schuerger et al. (7). Hypobaric (24 mbar), anoxic, CO₂-enriched atmospheric conditions were maintained within a vacuum desiccator to simulate a potentially habitable shallow martian subsurface microenvironment (Fig. 9). Here, CO₂-enriched refers to an atmosphere composed primarily of carbon dioxide; for reference, the present-day martian atmosphere consists of approximately 96% CO₂ (8). The system was flushed four times using a compressed gas cylinder containing 99.9% CO₂. Each flushing cycle consisted of evacuating the chamber to 24 mbar, repressurizing with CO₂, and evacuating again to 24 mbar. Five BD GasPak™ EZ Anaerobe sachets were placed within the desiccator to scavenge residual oxygen by converting it to CO₂. Chamber pressure was continuously monitored and maintained at or below 24 mbar using a vacuum pump controller (KNF Neuberger, Inc.).

**Figure 9.**
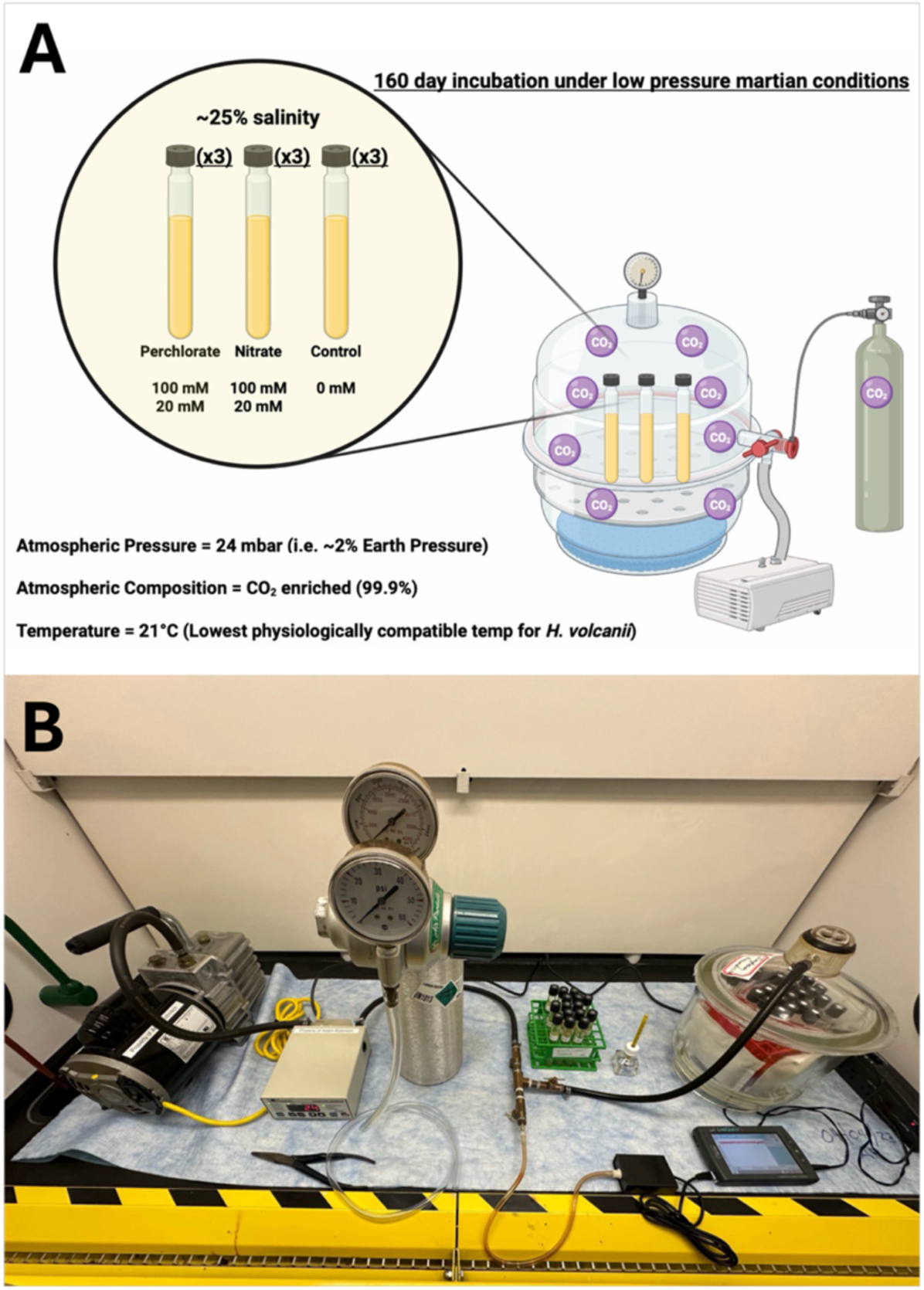
Incubation system used to incubate Haloferax volcanii cultures in hypersaline growth medium (225 g L⁻¹ NaCl) under defined subsurface martian conditions (99.9% CO₂, 24 mbar, 21 °C) for 160 days. (A) Schematic representation of the experimental setup showing triplicate cultures amended with perchlorate, nitrate, or no electron acceptor (control) maintained within a low-pressure CO₂-enriched atmosphere. (B) martian culture apparatus consisting of a vacuum pump, pressure controller, CO₂ supply, and sealed vacuum incubation chamber. Atmospheric pressure was regulated and maintained at no greater than 24 mbar throughout the experiment by the vacuum pump controller.

To permit pressure equilibration between the internal tube environment and the external low-pressure chamber while preventing microbial contamination during vacuum flushing, sterile 0.02 µm mixed cellulose ester membrane filters were installed in Hungate tube caps in place of butyl rubber stoppers. During method development, triplicate test tubes containing standard microbiological media (tryptic soy broth and lysogeny broth) were subjected to the full vacuum cycling protocol and subsequently incubated at 37 °C for 7 days to assess sterility. In addition, throughout all *H. volcanii* cultivation experiments, triplicate blank ATCC-974 media controls were included to monitor for contamination. At the conclusion of each experiment, 1 mL aliquots from blank media controls were stained with the nucleic acid dye 4′,6-diamidino-2-phenylindole (DAPI) to confirm the absence of microbial contamination. Briefly, 1 mL of sample was combined with 10 µL of a 10⁻³ M DAPI solution and 25 µL of a 10% formalin solution prepared in ATCC-974 salt solution. Samples were incubated for 30 min, filtered through black 0.22 µm membrane filters (Millipore GTBP02500), mounted on glass slides, and examined using epifluorescence microscopy. All contamination screening methods, including both culture-based and microscopic analyses, yielded negative results across all experiment series.

Water loss was monitored throughout the experiment because chamber pressure was near the vapor pressure of water at the incubation temperature (21°C). Following inoculation and placement of filter caps, individual test tubes were weighed immediately prior to placement in the vacuum desiccator. Final mass measurements were obtained at the conclusion of the incubation period, and water loss was calculated as the difference between initial and final mass. Culture volumes were restored to their initial values using 0.22 µm filter-sterilized buffer solutions prepared at the calculated final salt concentration of the media (i.e., ATCC-974 salt solution) to minimize osmotic disruption. To verify that mass measurements accurately reflected water loss and not scale drift, empty control tubes fitted with filter caps were included alongside experimental samples. Differences in mass for control tubes were within ±0.001 g, confirming reliable scale calibration throughout the experiment.

### Growth Measurements

Growth was indirectly assessed by measuring increases in optical density at 630 nm (OD_630_) of liquid broth cultures, used as an indication of increased cell density as described by Oren et al. (45). For both “martian” and “Earth control” cultures, 150 μL aliquots were collected in triplicate from each sample and transferred to a 96-well microtiter plate. Initial and endpoint optical density measurements were obtained to calculate changes in OD_630_. Measurements were taken spectrophotometrically using a BioTek ELx800 Absorbance Reader. For “Earth control” cultures incubated outside the Mars simulation chamber, optical density measurements were additionally collected once per week over the 160-day incubation period to enable semi-continuous monitoring of growth. These measurements were obtained by directly placing Hungate test tubes into the test tube holder of a Thermo Scientific™ SPECTRONIC™ 200 spectrophotometer. Semi-continuous measurements were not possible for martian cultures incubated under anoxic, low-pressure conditions, as opening the vacuum chamber during the incubation period would introduce oxygen and compromise experimental integrity. Consequently, only initial and endpoint OD_630_ values were recorded for “martian” cultures.

A viability assay was conducted for all cultures at the conclusion of the 160-day incubation period. 100 µl aliquots from all samples were transferred to 5 ml of fresh growth media in optimal, oxygenated growth conditions and incubated at 37 °C for 7 days, after which OD_630_ was measured. Cultures exhibiting an increase in OD_630_ during this recovery period were considered viable. Viability assays were performed in triplicate using standard ATCC-974 medium containing 125 g L⁻¹ NaCl as well as a modified ATCC-974 medium containing 225 g L⁻¹ NaCl. None of the growth media used in the viability assay included added nitrate or perchlorate.

For all treatments under anoxic conditions, tubes lacking an added electron acceptor served as negative controls to quantify growth attributable solely to dissolved oxygen. Growth on alternative electron acceptors was calculated as the change in OD (ΔOD_630_) relative to the final OD_630_ measured in the corresponding negative controls. To rule out microaerobic or abiotic oxyanion reduction, fully aerobic cultures containing nitrate or perchlorate and cell-free controls were incubated under identical chemical conditions. No oxyanion loss was detected in either condition, indicating that the oxyanion reduction observed in anoxic cultures was biologically mediated. All experiments were performed in triplicate sets.

### Electron acceptor measurements

As previously described in Robinson & McQuaig-Ulrich (43, 54) initial perchlorate concentrations were confirmed, and final concentrations were determined using perchlorate and nitrate selective ion-selective electrodes (Cole-Parmer Combination ISE) connected to an Oakton 450 pH/ISE meter. For analysis, 300 μL aliquots were withdrawn from each culture tube and transferred to a 10 mL beaker. Each sample was diluted 25-fold by adding 7.2 mL of deionized water. Subsequently, 150 μL of a 2 M ammonium sulfate ionic strength adjuster (ISA) was added to the diluted 7.5 mL sample, yielding the required 1:50 ISA-to-sample ratio. Measured concentrations were corrected for dilution to calculate the original ion concentrations in culture. For reference, a theoretical concentration of 0.1 M perchlorate (9945 mg L⁻¹) in culture would be measured as 397.8 mg L⁻¹ following dilution. Statistical analyses were performed using GraphPad Prism, by paired comparison of initial and final perchlorate concentrations for each biological replicate using two-tailed paired t-tests.

### Scanning Electron Microscopy

To examine possible changes in cell morphology under the tested conditions, all treatments were examined by scanning electron microscopy (SEM). Cultures were centrifuged, pellets collected, and treated for SEM. Cells and biofilm were prepared by using a procedure modified from Robinson & McQuaig-Ulrich (43). All steps involved used 1 mL of the described reagents or solutions. First, pellets were rinsed three times with ATCC 974 salt solution that was prepared at the final concentrations of the culture media at the end of the experiment. They were then fixed in a 2% solution of glutaraldehyde in 225 g L⁻¹ NaCl ATCC 974 salt solution at 4°C for 48 h.

The pellet was rinsed three times with 225 g L⁻¹ NaCl ATCC 974 salt solution. Following a dehydration series of 30, 50, 70, 90, 100, 100, 100% ethanol, 2:1 ethanol/hexamethyldsilazane, 1:1 ethanol/hexamethyldsilazane, and 100, 100, 100% hexamethyldsilazane. In each step, the pellet was incubated for 10 min. After the final dehydration step, the pellet was collected on a black 0.22 μm filter (Millipore GTBP02500) and dried in a desiccator overnight. The filter pad was then mounted on an SEM sample stub and sputter-coated with Au/Pt alloy. Samples were observed with a JEOL JSM6490 Scanning Electron Microscope.

### Raman Spectroscopy

Target peaks of interest for bacterioruberin (∼1505 cm^−1^, ∼1150 cm^−1^, and ∼1000 cm^−1^) were chosen as *H. volcanii* has a total carotenoid content of 82% bacterioruberin (51,60,61). Since β-carotene and bacterioruberin share similar Raman vibrational modes, we can use spectral peaks and intensities to determine *H. volcanii* growth. Samples were visually inspected with a 10x objective, allowing for features that indicated sufficient cell growth to be observed spectrally. Once in a viable location, Raman observations were conducted with acquisition and integration times ranging from 2-7 minutes, allowing for weaker intensity peaks to be measured, if present. Spectra were taken and averaged within a given micrometer region of the observed region. Raw data was processed and spectra were smoothed to illustrate the carotenoid peaks present, or lack thereof.

## Acknowledgments

The authors thank Dr. Jamie Foster, Dr. Andy Cannons, Dr. Mya Breitbart, Amanda Garces, Dr. B. Jake Cha, Dr. Basem Jaber, Makenzie Kerr, John Foster, Dr. Erin Goergen, Dr. Natavia Middleton, Justin Hubsmith, and Dr. Corrado Caslini for their guidance, support, and helpful discussions.

This work has been supported in part by the Lisa Muma Weitz Microscopy Core Laboratory at University of South Florida College of Medicine. We would like to especially thank Amanda Garces for her expert assistance with SEM imaging and acquisition of the micrographs.

This research received no specific grant from any funding agency in the public, commercial, or not-for-profit sectors.

